# Genetic Mapping of SNP Markers and Candidate Genes Associated with Day-Neutral Flowering in *Cannabis sativa* L

**DOI:** 10.1101/2023.04.17.537043

**Authors:** Andrea R. Garfinkel, Dustin G. Wilkerson, Hsuan Chen, Lawrence B. Smart, Brendan M. Rojas, Brooke A. Getty, Todd P. Michael, Seth Crawford

## Abstract

Hemp (*Cannabis sativa* L.) is a primarily short-day crop grown for grain, fiber, and secondary metabolites. Although terminal flowering of most non-domesticated hemp is regarded as photoperiod sensitive, limited germplasm is available for the development of day-neutral hemp. Day-neutral plants experience flower maturation independently of the short-day photoperiod cue which is typically triggered by photoperiods of less than 14 hours. The day-neutral trait is the subject of increased commercial interest for the purpose of breeding varieties with accelerated flowering time or cultivars that do not require labor- and cost-intensive light deprivation production systems. Genetic markers for the photoperiodic response would help breeders make early selection in the design of suitable cultivars for specific environments and cultivation calendars. Limited refereed information has been produced regarding the genetic regulation of photoperiodism in *C. sativa*. A population of 318 F_2_ individuals segregating for day-neutral flowering was developed, phenotyped, and genotyped. Genome-wide association analysis identified markers associated with the day-neutral trait indicating that a single recessive gene controlling photoperiod sensitivity is located within a large region of Chromosome 1. Flanking region sequence data of linked SNP markers was used in the development and validation of a TaqMan-based qPCR assay for the day-neutral trait. A genetic linkage map was produced, and QTL mapping identified two additional markers on Chromosome 1. Candidate genes, *TARGET OF EARLY ACTIVATION TAGGED (TOE*)/*APETALA2* (*AP2*), and *PSEUDO-RESPONSE REGULATOR 3* (*PRR3*), may work together to impact phase transition and photoperiodic flowering respectively and have key domains that are disrupted in day-neutral plants.

The day-neutral flowering (a.k.a. “autoflower”) phenotype is a valuable and understudied trait in *Cannabis sativa*

The phenotype may result from a conserved linkage block of phase transition and photoperiodic flowering genes

A segregating population was developed, genome wide analyses identified markers linked to the day-neutral trait

Correlated markers were mapped to Chromosome 1 across a sizable region that appears to be largely non-recombining

## INTRODUCTION

*Cannabis sativa* L. is an annual row crop grown for protein- and oil-rich grain, multiple industrial usage fibers, and for the secondary metabolites produced by the plant (referred to as “cannabinoids”) for medicinal and recreational use (Henry et al., 2020; McPartland, 2018). In recent years, scientific interest in the development of hemp cultivars for medicinal use has increased (Lewis et al., 2018; Toth et al., 2020) due to the legalization of *C. sativa* with total Δ^9^- tetrahydrocannabinol (THC) < 0.3% in the United States and elsewhere, and the subsequent broadening of markets for cannabinoids (Walia, 2019). Cultivation of *C. sativa* for cannabinoid production is focused on the harvest and yield of female flowers. Flower bracts, especially those on female plants, contain high densities of structures known as glandular trichomes within which cannabinoids are produced (Mahlberg and Kim, 2004). At least 150 cannabinoids have been discovered with potentially different medicinal or psychoactive functions (Hanuš et al., 2016), of which cannabidiol (CBD), THC, and cannabigerol (CBG) are produced in the greatest quantities.

The transition from vegetative growth to flowering in *C. sativa* has historically been considered to be regulated by photoperiod. However, recent investigations argue that development of small solitary flowers adjacent to the leaf axils of *C. sativa* occurs regardless of photoperiod (Spitzer-Rimon et al. 2019). The authors of this study furthermore suggest that it is in fact the flower *maturation* (as opposed to *initiation*) and development of terminal inflorescences processes that are regulated by photoperiodic cues (Spitzer-Rimon et al. 2019). It is with this research in mind that we consider our further investigations into the photoperiodic response of *C. sativa* and classify the terms “photoperiod sensitive” and “day-neutral” as related to the development and maturation of terminal inflorescences as an observable phenotype and binary trait. A recent study by Toth et al. (2022) confirmed that the the photoperiod sensitivity of *C. sativa* is likely regulated by a single recessive gene.

Most *C. sativa* plants behave as short-day plants, requiring a minimum dark period for inflorescence development (Hall et al., 2012). Studies have shown that most *C. sativa* plants will transition to flower maturation (classically considered to be the transition from the “vegetative” to “flowering” stage) during photoperiods less than 14-16 h, depending on cultivar (Hall et al., 2012, Zhang et al., 2021). Although most *C. sativa* plants are photoperiod sensitive, some genotypes display a day-neutral trait, often referred to colloquially as “autoflowering.” Day-neutral *C. sativa* plants develop terminal inflorescences regardless of photoperiod (Zhang et al, 2021), sometimes in response to plant stressors such as excess heat or being root-bound. The origin of the day-neutral background in cannabis is not clearly resolved, in part due to confusing and shifting historical taxonomic classifications within the genus. However, those with this trait are sometimes ascribed to the “Ruderalis” type, as potentially originating from a taxonomically unresolved subspecies or variant of *C. sativa*, once referred to as *C. ruderalis* (or *C. sativa* var. *ruderalis*) (McPartland, 2018). The validity of this species or subspecies distinction is tenuous and the subject of debate; nonetheless, it is worth acknowledging this distinction in order to place this trait within the *Cannabis* species literature. Regardless of the systematics, day-neutral genotypes are typically considered to be from northern Eurasia (Small, 2018).

“Autoflowering” has become an important buzzword in *C. sativa* breeding as its desirability as a trait continues to increase. Not only can the day-neutral background help in breeding efforts to reduce time between generations, but this trait has become important for farmers as well. Terminal flower development in short-day, photoperiod-sensitive cultivars is dependent on daylength; therefore, time to flower varies among growing regions at different latitudes, with the day-neutral plants suspected to be specially adapted to survival at the northernmost latitudes which experience long days late into the growing season (Small, 2018). In some cases, the ability to override the photoperiod-sensitivity of *C. sativa* is desirable at lower latitudes as well. In more southern locations with warmer climates and long growing seasons, the day-neutral trait can allow for two consecutive harvests in one season. Furthermore, day-neutral cultivars can be produced year-round in greenhouse facilities without the need for costly or labor-intensive light deprivation capabilities. Despite the increasing interest in day-neutral cultivars, few studies have investigated the effect of this trait on cannabinoid content, yield, or other agronomic traits; however, there is some indication that the pattern of accumulation of cannabinoids in day-neutral cultivars is different than in photoperiod-sensitive cultivars (Yang et al., 2020).

Selection for plants with the day-neutral phenotype in the breeding process is complicated. Day-neutral plants are phenotypically indistinguishable from photoperiod sensitive plants prior to terminal flowering; therefore, phenotyping requires growing under long days and waiting for inflorescence development and maturation. At this point, plants must be quickly utilized for breeding because day-neutral plants will begin to senesce and die following flowering, making them very difficult to propagate clonally using vegetative cuttings or in micropropagation (Piunno et al., 2019). In this way, there is a short, one-time window for making selection and crosses, which, if missed, delays breeding efforts.

Due to the challenges of breeding with day-neutral plants, molecular markers to identify individuals with this trait prior to flowering is desirable. The purpose of this study was to utilize a newly developed, *Cannabis*-specific SNP array (also known as a “SNP chip”) in mapping the locus/loci controlling the day-neutral trait in a segregating F_2_ population of *C. sativa* and in the identification of linked sequence polymorphisms. Our goal was to further utilize these polymorphisms to develop a rapid-screening qPCR assay for use in the early selection of day-neutral plants in the breeding process. We report herein the approximate genomic location of the markers developed, elucidate potentially unique patterns of recombination along Chromosome 1 of *C. sativa* and show that candidate genes, *TARGET OF EARLY ACTIVATION TAGGED (TOE*)/*APETALA2* (*AP2*), and *PSEUDO-RESPONSE REGULATOR 3* (*PRR3*), are linked over evolutionary time from basal angiosperm to monocots and both have disrupted conserved domains in day-neutral plants.

## MATERIALS AND METHODS

### Development of the segregating population

A biparental F_2_ population of 318 individuals segregating for photoperiod sensitivity was developed. The F_1_ plant was a hybrid from the seed parent, *C. sativa* ‘HO40’, a photoperiod-sensitive plant (additional description of the plant ‘HO40’ can be found in Garfinkel et al. 2021), and the pollen plant, *C. sativa* ‘ERB’, a CBD-dominant, day-neutral plant. Both *C. sativa* ‘HO40’ and *C. sativa* ‘ERB’ are dioecious, female inbred lines maintained by Oregon CBD (Independence, Oregon) and are all stable in their photoperiodic response. One single female F_1_ plant was self-pollinated to create the F*2* mapping population in a 60 x 60 x 120 cm^3^ growth tent (TopoLite, Amazon) equipped with a 600W LED light (Phlizon, Guangdong, China) and inline fan (vtx600, Terrebonne, Québec, Canada) and HEPA filter (Phat IGS6HEPA, Hydrofarm) for ventilation and to avoid external pollen contamination.

In all crosses, a modified silver thiosulfate (STS) treatment (Ram and Sett, 1982) was used to stimulate male flower development on genetically female plants for self-pollination and inbred line maintenance (de Meijer et al., 2003). The STS buffer was applied three times, once every five days, with the first application simultaneously coinciding with a reduction in photoperiod to 12 h/12 h light/dark cycle for flower induction. In the case of the initial hybrid cross, the entire pollen parent was treated, whereas in order to self the F_1_ plant, only a single branch was treated with STS to stimulate male flower development. Pollen from male flowers was then used to hand pollinate the female flowers on the rest of the plant that had not been converted. Mature F_2_ seeds from the F_1_ plant were collected and sown in seedling trays. F_2_ seedlings were transplanted two weeks after sowing into 26.5-L pots filled with organic soil medium (Organic Valley with Worm Castings, Rexius, Eugene, Oregon) and moved into a research greenhouse at Oregon CBD (44°53’01.84” N, 123°13’51.81” W) where they were grown under continuous supplemental light for 140 days for phenotyping.

### Phenotyping and inheritance of phenotype

The 318 F_2_ progeny for phenotyping were grown under continuous light to distinguish photoperiod sensitive and day-neutral individuals. Given that both day-neutral and photoperiod sensitive plants will develop solitary flowers at sexual maturity (Spitzer-Rimon et al., 2019), the photoperiodic response was assessed by observing the presence or absence of terminal inflorescence development under constant light conditions. Plants which did not develop terminal inflorescences were recorded as photoperiod-sensitive, whereas those with extensive flowering were recorded as day-neutral. No plants were observed as having an intermediate phenotype.

The segregation ratio of day-neutral and photoperiod-sensitive plants was then examined by goodness-of-fit chi-squared (χ^2^) tests. The expected segregation ratio of 1 day-neutral:3 photoperiod insensitive for a single recessive gene was used as the null hypothesis.

Seventeen additional cultivars unrelated to the parents used in the crossing population, eight day-neutral and nine photoperiod-sensitive were grown under the same cultural conditions as above (Supplemental Table S1). These plants were used to validate if the qPCR markers that we developed were predictive of the day-neutral trait outside of our genotyped population

### DNA extractions and genotyping

Of the 318 individuals, 192 plants, including 62 day-neutral plants and 130 photoperiod-sensitive plants, were randomly selected for genotyping and linkage map construction. DNA was extracted using a Quick-DNA Plant/Seed 96 Kit (Zymo Research, Irvine, California) according to the manufacturer’s instructions. DNA was sent for genotyping on the CannSNP90 SNP array (Eurofins BioDiagnostics, Grand Forks, North Dakota & Medicinal Genomics, Beverly, Massachusetts), which contained over 89,000 markers across the *C. sativa* genome. DNA from the additional unrelated 17 individuals used for validation of a qPCR assay were also extracted using a Quick-DNA Plant/Seed Miniprep Kit, except that samples were initially frozen in liquid nitrogen and homogenized.

### Marker-trait association analysis

A total of 75,081 SNPs were successfully genotyped by the CannSNP90 SNP array. After sorting and trimming the genotype dataset from the CannSNP90 array to eliminate markers that had ≥ 30% missing data or were ≤ 5% polymorphic, 16,272 markers were included in the genome-wide association (GWAS) analysis. Principal component (PCA) and marker-trait association analyses were performed using GAPIT (Lipka et al., 2012). PCA = 8 and several statistical models, including multivariate linear models (MLM), multiple locus mixed linear model (MLMM), and compression MLM (CMLM), were used for marker discovery. Potential markers from different models were compared and selected using QQ-plots, P value, Bonferroni adjusted P values, and FDR adjusted P values from the GAPIT analysis (Figure 1).

**Figure 1.**
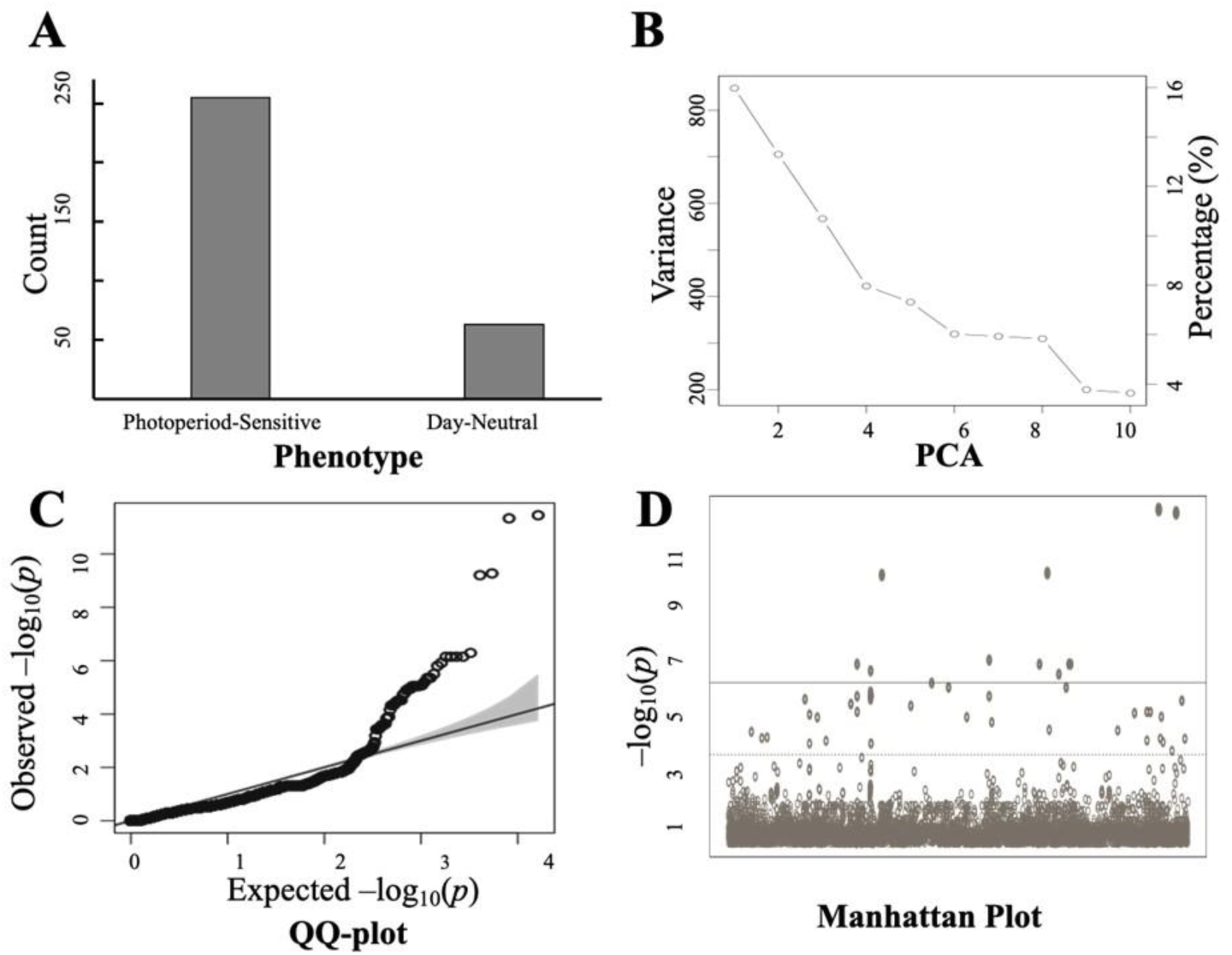
Phenotypic segregation of the day-neutral trait in *Cannabis sativa* and marker-trait association results produced using GAPIT (Lipka et al., 2012). Flowering phenotype of 318 non-selected F_2_ individuals from a ‘ERB’ (day-neutral) x ‘HO40’ (photoperiod sensitive) cross. (B-D) A total of 192 F_2_ plants, including 62 day-neutral plants and 130 photoperiod-sensitive plants, were analyzed in GAPIT using 16,272 loci from the CannSNP90 chip array. (B) PCA eigenvalues from 1 to 10, (C) QQ-plot analysis, and (D) the Manhattan plot produced from the GAPIT analysis with the Bonferroni correction threshold (solid line) and the FDR threshold (dash line) of α = 0.05.

### PCR/qPCR-based marker development

Flanking sequences on either side of the SNPs included in the array were provided by Medicinal Genomics. These were used to BLAST whole-genome sequences of ‘ERB’ and ‘HO40’ (or closely related individuals) using Geneious Prime (v. 2020.2.1) in order to identify additional variation in the regions of interest for primer and probe development. Several additional SNPs were located in the region, and these data were used to design a custom TaqMan® SNP Genotyping Assay using the Custom TaqMan® Assay Design Tool (Thermo Fisher Scientific, Waltham, Massachusetts) to detect the SNP used in the chip or another variable region within 100 bp of the originally detected SNP or indel locus (see results). Primers for end- point PCR were also developed (primer pairs SNP262_1F/SNP262_1R and SNP504_1F/SNP_504_1R) to amplify and sequence a 250-350 bp region surrounding the SNP in some of the F2 progeny. Sequenced fragments were used to validate the calls of the SNP chip and identify controls for use in qPCR SNP genotyping assays. All primers and probes are listed in Table 1. Flanking region sequences of ‘ERB’ with the SNP targeted in qPCR analyses identified are available as Supplementary Material.

**Table 1.**
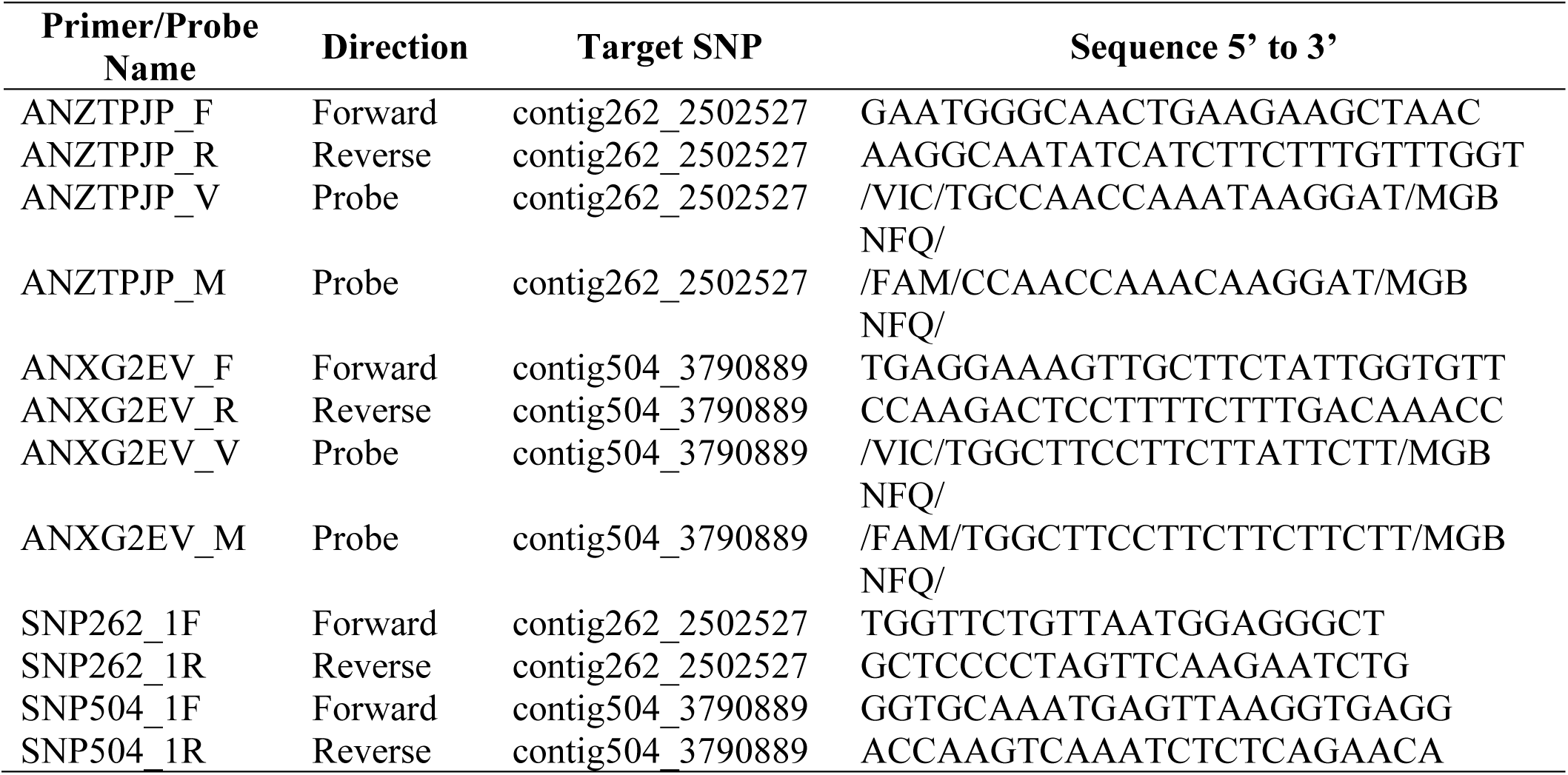
Primers and probes used to amplify and/or target regions associated with day-neutral flowering in Cannabis sativa.

End-point PCRs were performed in 20 µL reactions containing: 1x Platinum II Hot-Start Master Mix (Invitrogen, Carlsbad, California), 220 nM of each primer, water, and variable amounts of DNA (not quantified). Cycling conditions were: 94°C for 2 min followed by 35 cycles of 98°C for 5 sec and 60°C for 15 sec. PCR products were visualized on a FlashGel™ system (Lonza, Basel, Switzerland), cleaned up using ExoSAP-IT Express (Applied Biosystems, Foster City, California), and sent to sequencing using the same primers used in PCR (GENEWIZ, South Plainfield, New Jersey).

qPCR analyses to detect the SNPs were performed in 10 µL reactions using the TaqPath ProAmp Master Mix (Thermo Fisher Scientific) according to the manufacturer’s recommendations for their custom designed SNP genotyping assays. Fast genotyping run conditions were used on a QuantStudio 5 qPCR thermal cycler (Applied Biosystems). The QuantStudio 5 software was used to make automatic genotyping calls using three progeny identified during Sanger sequencing as controls (two homozygous controls and one heterozygous control). All automatic calls were visually scored for accuracy following the run.

### QTL mapping

All linkage and QTL mapping in this study was performed in R (R Core Team, 2022). Formation of the linkage map began with the same 75,081 markers that were successfully genotyped from the CannSNP90 SNP array. Of these, 55,579 markers were over 98 percent monomorphic in this population and removed. An additional filter isolating markers with call rates greater than 70 percent resulted in 16,200 polymorphic markers. These markers were further narrowed down using multiple sequence runs of the parents and F_1_, allowing for the classification of marker types based on consensus parental genotypes and the isolation of 9,746 intercross markers in this population. Linkage map construction and QTL mapping using these intercross markers was facilitated by a combination of R/qtl (Broman et l., 2003) and ASMap (Taylor and Butler, 2017). Following removal of co-located and extremely distorted markers, 7,256 markers were used to define the initial linkage groups using the ‘mstmap’ function (p.value = 1e-25, objective.fun = “ML”, dist.fun = “kosambi”, bychr = FALSE). This function clustered the markers into linkage groups and determined the marker order. Any marker that failed to cluster was removed. Marker order and linkage map stability of the final linkage map, totalling 1,103 markers across 10 linkage groups, were visually assessed using the functions ‘heatMap’, ‘profileMark’, and ‘plotMap’.

To improve interpretability with the hemp research community’s standard chromosome numbering, each marker in the linkage map’s 1 kb flanking sequence (provided by Medicinal Genomics based on ‘Jamaican Lion’ genome coordinates) was nucleotide blasted (Comacho et al., 2009) using default parameters to the ‘CBDRx-CS10’ reference genome, available on NCBI (https://www.ncbi.nlm.nih.gov/assembly/GCF_900626175.2). The ‘best’ BLASTn results for each marker were determined using the highest bitscore, based on overall alignment quality.

Linkage groups were cross-validated into representative chromosomes by identifying the most likely ‘CBDRx-CS10’ placement of ‘Jamaican Lion’ contigs by majority placement of their individually aligned markers. This allowed for the linkage groups to be assigned to their respective ‘CBDRx-CS10’ reference chromosomes.

The autoflower phenotype data described above was used in detecting QTL in the linkage map. QTL mapping was performed using functions from R/qtl. First, genotypic probabilities were calculated using ‘calc.genoprob’ (step = 5, map.function = “kosambi”) followed by ‘scanone’ (model = “binary”, method = “em”) for QTL mapping. The permuted 5% genome-wide significance threshold was determined by using ‘scanone’ again with the addition of n.perm = 1000. Lastly, ‘refineqtl’ and ‘fitqtl’ (method = “hk”, model = “binary”) followed by ‘lodint’, determined the most likely position of the peak LOD and established the 1.5 LOD support interval.

### Candidate gene analysis

Since the day-neutral trait is recessive, known day-neutral recessive plants (those that contained one copy of the day-neutral chromosome and one copy of the day-sensitive chromosome as a result of a cross between homozygous parents of representing each phenotype) were selected for further analysis: ‘ERB’ (day-neutral) x ‘HO40’ (day-sensitive); ‘SSxHO40’ (day-sensitive) x ‘SSA’ (day-neutral); ‘WeddingCakexFB30’ (day-neutral) x ‘HO40’ (day- sensitive). High molecular weight (HMW) DNA was extracted from the day-neutral recessive plants, sequenced on the Pacific Biosciences (PacBio) Sequel II sequencer, assembled with HiFiasm, and protein coding genes were predicted (A. Garfinkel, T. Michael, and S. Crawford, *in preparation*). The protein coding sequences for TOE/AP2 and PRR3 were aligned across the three day-neutral recessive plants with MUSCLE (**MU**ltiple **S**equence **C**omparison by **L**og- **E**xpectation) algorithm using default settings in the program MEGAX (v10.1.8). The evolutionary relationship between TOE/AP2 and PRR3 was explored using the full network synteny based analysis of 123 high quality plant genomes spanning basal angiosperm to monocots (Zhao et al., 2021). All syntenic blocks across the 123 plant genomes were selected if they contained the orthogroup for both TOE/AP2 and PRR3. The syntenic blocks were plotted in a selected number of plant genomes highlighting the linked genes using MCscan Python version (JCVI utility libraries v1.3.3).

## RESULTS AND DISCUSSION

### Phenotype segregation

Of the phenotyped 318 F_2_ individuals, 255 plants were photoperiod-sensitive, and 63 plants were day-neutral. Using these data, the expected 1:3 segregation ratio for a recessive allele at a single locus was rejected by the *χ*^2^ goodness-of-fit test (*χ*^2^= 4.566, df = 1, *P* value = 0.032, α=0.05). We can only speculate as to the reason for this segregation distortion; it is possible that certain allelic combinations were lethal, less vigorous, or there was unknown epistasis. We have sometimes observed higher rates of, or harder-to-break dormancy, as well as delayed germination in plants with the day-neutral phenotype (S. Crawford, A. Garfinkel, and B. Getty, unpublished data). It is possible that the segregation ratios were skewed due to differences in germination rates at the time of phenotyping the crossing population. A study done by Beutler and Der Marderosian (1978) reported a similar delay in germination of seeds in crosses between day-neutral and photoperiod sensitive plants; the authors speculate this trait is a carryover from the “*C. ruderalis*,” or day-neutral, parent. Although cultivars containing the day-neutral gene have been used in segregation studies since at least the 1970s (Beutler and Der Marderosian, 1978), this is the one of the first studies, that we are aware of, to report the inheritance model of the day-neutral trait by creating a segregating population. Beutler and Der Marderoian created a day-neutral/photoperiod sensitive F_1_ population but was not reported to segregate for photoperiod sensitivity. Toth et al. (2022) recently described an F_2_ pedigree similar to the current study. Interestingly, Beutler and Der Marderosian (1978) reported that in their day-neutral/photoperiod sensitive cross, using the day-neutral plant as the seed-bearing parent resulted in very few seeds and progeny with poor vigor. Our recent pangenome analysis of day-neutral vs. day-sensitive plants revealed inverted repeats that may impact recombination on Chromosome 1 (*in preparation*). Additional data regarding the inheritance of the day-neutral trait, possible lethal combinations, any asymmetric crossing compatibility, and/or epistatic effects in other crossing populations is warranted. Although the segregation data we collected from this single population do not fit the 1:3 segregation ratio, the genotyping results discussed below provide ample evidence to support the single gene model with a recessive locus, suggesting that phenotyping of additional pedigrees is warranted to provide confirmation.

### Marker-trait association and qPCR assay development

Several statistical models were evaluated using GAPIT, including MLM, MLMM, and CMLM. All models produced similar results insofar as the same candidate markers were selected by multiple models, but with varying yet still significant *P* values. The MLMM model (Segura et al., 2012) from GAPIT was ultimately used to select markers, therefore, the results of the MLMM model are presented. We further utilized the Quantile-Quantile plot (QQ-plot) and Manhattan plot with a Bonferroni threshold to examine the fitness of the MLMM model and confidence levels of the selected markers (Figure 1). Without using marker location information, GAPIT found two candidate markers, contig504_3790889 and contig262_2502527, that have the strongest associations with the day-neutral trait with *P* values of 0 and 3.55 x 10^-12^, respectively (Figure 1). The FDR adjusted *P* values of two candidate markers, contig504_3790889 and contig262_2502527, were 0 and 2.52 x 10^-8^, respectively (Figure 1). Subsequent analysis of the contig504_3790889 marker showed that this locus segregated perfectly with the day-neutral versus photoperiod sensitive phenotype.

According to the CannSNP90 genotyping report, the contig504_3790889 marker is a SNP locus with A/C alleles and the contig262_2502527 marker is placed at an indel. While we predicted to see three genotypes in the F_2_ population (AA, AC, and CC) at contig504_3790889, only genotypes AC and CC were recorded (Table 2). Only 184 of the 192 sequenced plants were successfully genotyped at this locus. Plants with the AC genotype were all day-neutral plants (n = 54), whereas plants with the CC genotype were all photoperiod-sensitive plants (n = 130) predicting the day-neutral trait in this population without error (Figure 2).

**Figure 2.**
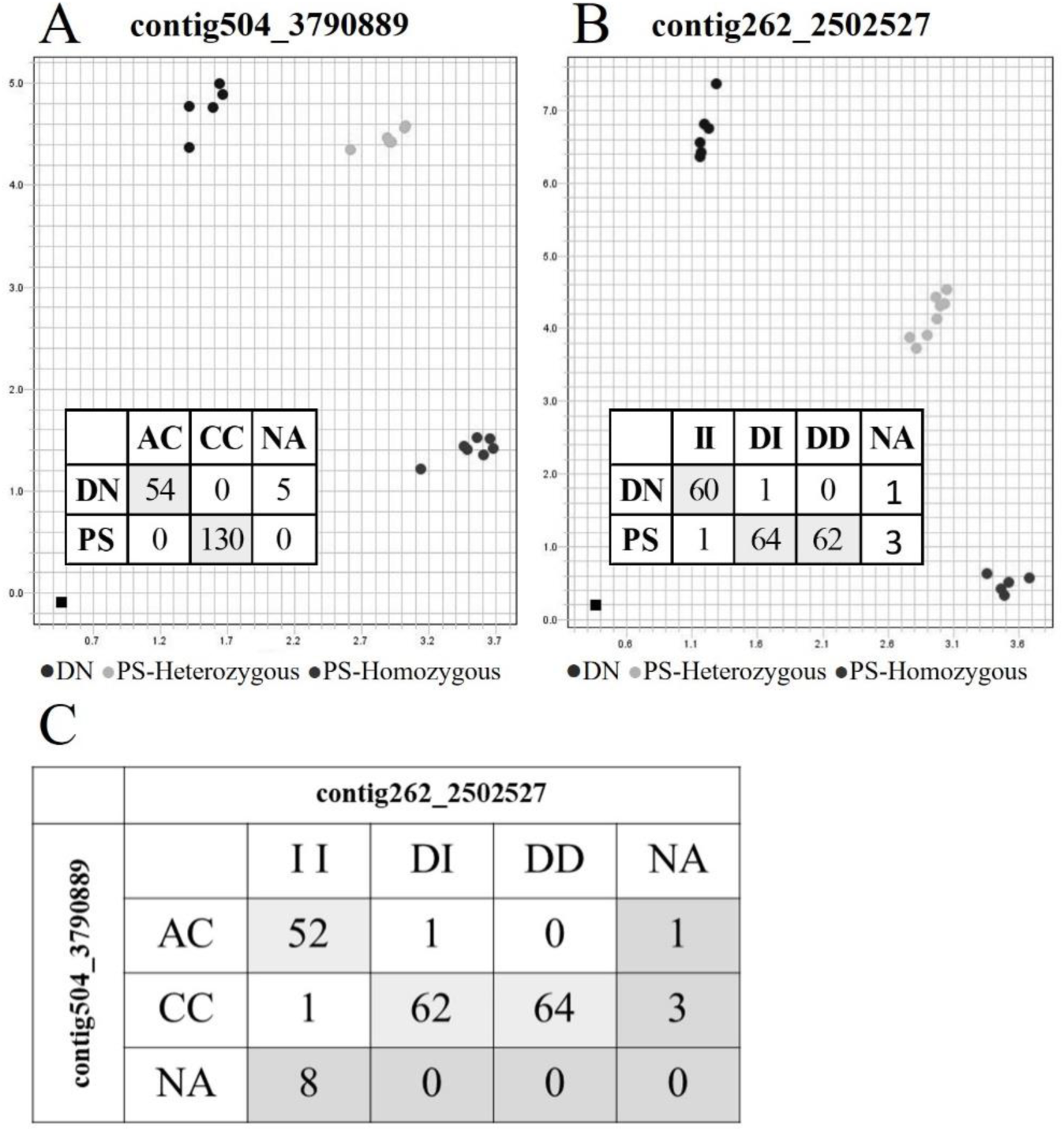
Allelic segregation of the two SNPs associated with the day-neutral trait in *Cannabis sativa*. (A) The table indicates the two genotypes identified by the CannSNP90 array at the contig504_3790889 locus in an F_2_ population consisting of both day-neutral (DN) and photoperiod sensitive (PS) individuals; a qPCR assay (SNP_504) was developed, validated, and identified three genotypes at this locus associated with flowering behavior in the segregating population as seen in the allelic discrimination plot. (B) The table indicates genotypes identified in the same population at locus contig262_2502527 by the CannSNP90 and the allelic discrimination plot shows validation of a qPCR assay (SNP_262). “NA” indicates the number of individuals for which genotyping calls were not made at a given locus. (C) shows the relationship between genotype calls by the CannSNP90 at the two loci associated with the day-neutral phenotype.

**Table 2.**
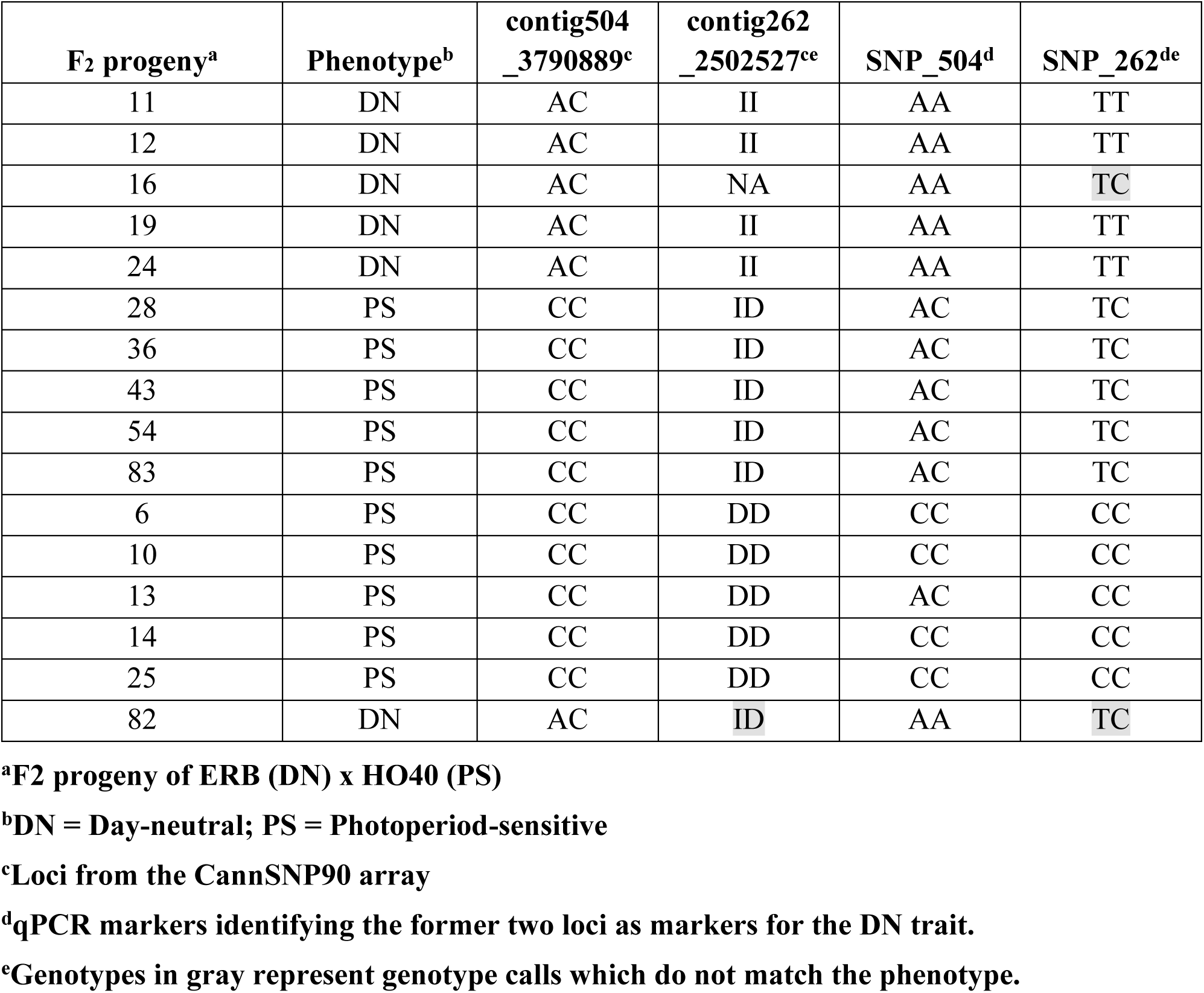
Genotypes at the two day-neutral associated loci from the CannSNP90 array, contig504_3790889, contig262_2502527, and the two loci as converted into qPCR assays, SNP_504 and SNP_262, for the 16 F_2_ individuals.

| F<sub>2</sub> progeny<sup>a</sup> | Phenotype<sup>b</sup> | contig504_3790889<sup>c</sup> | contig262_2502527<sup>ce</sup> | SNP_504<sup>d</sup> | SNP_262<sup>de</sup> |
| --- | --- | --- | --- | --- | --- |
| 11 | DN | AC | II | AA | TT |
| 12 | DN | AC | II | AA | TT |
| 16 | DN | AC | NA | AA | TC |
| 19 | DN | AC | II | AA | TT |
| 24 | DN | AC | II | AA | TT |
| 28 | PS | CC | ID | AC | TC |
| 36 | PS | CC | ID | AC | TC |
| 43 | PS | CC | ID | AC | TC |
| 54 | PS | CC | ID | AC | TC |
| 83 | PS | CC | ID | AC | TC |
| 6 | PS | CC | DD | CC | CC |
| 10 | PS | CC | DD | CC | CC |
| 13 | PS | CC | DD | AC | CC |
| 14 | PS | CC | DD | CC | CC |
| 25 | PS | CC | DD | CC | CC |
| 82 | DN | AC | ID | AA | TC |

<sup>a</sup>F<sub>2</sub> progeny of ERB (DN) x HO40 (PS)
<sup>b</sup>DN = Day-neutral; PS = Photoperiod-sensitive
<sup>c</sup>Loci from the CannSNP90 array
<sup>d</sup>qPCR markers identifying the former two loci as markers for the DN trait.
<sup>e</sup>Genotypes in gray represent genotype calls which do not match the phenotype.

Sequence data from ‘HO40’ and ‘ERB’ (Supplementary Data) and from selected F_2_ progeny generated with the SNP504_1F/SNP_504_1R primer pair indicated that the flanking region surrounding the contig504_3790889 SNP contains an imperfect, trinucleotide microsatellite with a CTT motif, with the target SNP located in a 3bp insertion towards the 5’ end of the repeats. This motif is repeated 7 times in ‘HO40’ and 4 times in ‘ERB’ prior to the targeted SNP (in reverse complement, GAA, in Supplementary Data); the motif was repeated 5 times in the flanking region sequence data provided by Medicinal Genomics. We do not know the probe sequence used in the CannSNP90 array to make definite conclusions, but it is possible that depending on probe design, some genotyping errors could be expected due to mis-annealing along this highly repetitive region causing the unexpected allele calls described above.

With the strong apparent linkage to the day-neutral locus as determined by the *P* value of 0, the location information of this marker demonstrated the greatest potential to be used in the design of qPCR-based assay. The variability at this locus elucidated by the sequence data (included as Supplementary Data) allowed for the development of qPCR SNP assay “SNP_504” (primers and probes beginning with ANXG2EV, Table 1). SNP assay SNP_504 helped resolve and correct genotype calls made by the CannSNP90 array; day-neutral plants were found to have the genotype AA, and both the AC and CC genotypes were detected in the photoperiod-sensitive plants (Table 1 and Figure 2). The qPCR SNP assay SNP_504 correctly predicted the photoperiodic response of 16 of the phenotyped and genotyped F_2_ progeny (Table 1 and Figure 2). However, when the qPCR SNP_504 assay was used on the 17 unrelated *C. sativa* cultivars (Supplemental Table S1), all of the plants genotyped regardless of actual phenotype were scored as the AA genotype, associated in our cross with the day-neutral trait (Supplemental Table S1). These results indicate that this assay will not likely be widely applicable in other crosses due to variability at this locus between our F_2_ population and other, non-related cultivars. Nonetheless, the data we have provided herein, specifically, the flanking sequence data (Supplemental Material) and primer sequences for Sanger sequencing this locus (Table 1), could be useful in designing a qPCR assay for specific crossing populations on a case-by-case basis, assuming the same locus is responsible for the day-neutral phenotype. The putatively universal markers developed by Toth et al. (2022) were not publicly available at the time of this study and were not tested in the current crossing population or 17 unrelated cultivars.

The other discovered marker, contig262_2502527, also explains the day-neutral trait with 99% correct prediction. At the contig262_2502527 locus, 188 F_2_ plants, including 61 day-neutral plants and 127 photoperiod-sensitive plants, were successfully genotyped using the CannSNP90 array (Figure 2). Unlike the data at the contig504_3790889 where only two genotypes were detected, the CannSNP90 array call data from locus contig262_2502527 matched the expectation of the population structure and contained three genotypes: II (homozygous for the insertion), ID (heterozygous insertion/deletion), and DD (homozygous for the deletion). Of the 192 individuals genotyped, a total of 61 plants were scored as II at this locus; of these 61, 60 plants were day-neutral (98.4%), and only one plant was photoperiod-sensitive (1.6%), suggesting that the insertion is strongly associated with the day-neutral trait in our population, although some infrequent crossing over events occur. Of the other 127 plants that genotyped either ID or DD at this locus, one plant was day-neutral (1.5%), and 126 plants were photoperiod-sensitive (98.5%). These results suggest that while not as strongly linked to the day-neutral trait as the contig504_3790889 locus, the contig262_2502527 locus is also reliable marker for predicting the day-neutral trait.

Due to the location of the indel used on the CannSNP90 array and the sequence variability in the flanking regions of this locus as identified by sequence data, a qPCR assay to target the indel was not feasible. Therefore, a nearby SNP was targeted for assay development. SNP qPCR assay SNP_262 (primers and probes beginning with ANZTPJP, Table 1 and Figure 2) was designed to target a T/C SNP with the TT genotype corresponding to the day-neutral trait. The qPCR genotyping calls agreed 100% with the data generated by and predicted phenotype as reliably as the CannSNP90 array, with two calls of the 16 selected individuals for qPCR SNP assay validation failing to accurately correspond with phenotype (Table 2); the same incorrect calls made by the qPCR assay were also made by the CannSNP90 array, suggesting a recombination event in between locus contig262_2502527 and the gene/locus responsible for the day-neutral trait. Like qPCR assay SNP_504, SNP assay SNP_262 also failed to accurately predict phenotype in the additional 17 unrelated individuals tested (Supplemental Table S1). Therefore, this marker is unlikely to be widely applicable for use in breeding and selection for the day-neutral phenotype. Nonetheless, the data presented herein should provide a starting point for re-design of successful assays for other *C. sativa* populations in the case the same genomic region is associated with the day-neutral trait in those crosses. Toth et al. suggest that it is possible there are multiple loci regulating the day-neutral trait; our results in an unrelated crossing population to the one used by Toth et al. suggests the same locus may be responsible for the day-neutral trait in both studies.

The two markers, contig504_3790889 and contig262_2502527, appear to be strongly linked to one another (Figure 2). A total of 144 of the 192 plants genotyped yielded information for both contig504_3790889 and contig262_2502527; the genotypes of these two markers were strongly correlated (*χ*2 = 135.64, df = 2, *P* value = 3.51 x 10^-30^). At the contig504_3790889 locus, 53 plants were genotyped AC; of those 53 plants, 52 and 1 of them, respectively, had the II and DI genotypes at contig262_2502527 (Figure 2). Similarly, 127 plants were genotyped CC at the contig504_3790889 locus; 126 of them were called as DD or DI at contig262_2502527 (Figure 2). Only one plant that was genotyped CC in the contig504_3790889 locus was genotyped II in contig262_2502527 (Figure 2).

We earlier rejected the single recessive gene model for the day-neutral response; however, a marker at a single SNP locus was successful in predicting the day-neutral phenotype. Given these results, it is still feasible that this model is true and segregation ratios were skewed as a result of unequal germination or inverted repeats impacting recombination. This hypothesis warrants further exploration as recent studies have also shown that the day-neutral trait is likely governed by a single recessive gene in the same location (Toth et al., in review).

### Genetic linkage and QTL mapping

A total of 1103 genetic markers from the SNP array were divided into 10 linkage groups, corresponding with the 10 chromosomes of *C. sativa* (nine autosomes and a sex chromosome). The resultant genetic map is shown in Figure 3 and summary statistics for the linkage map are shown in Table 3.

**Figure 3.**
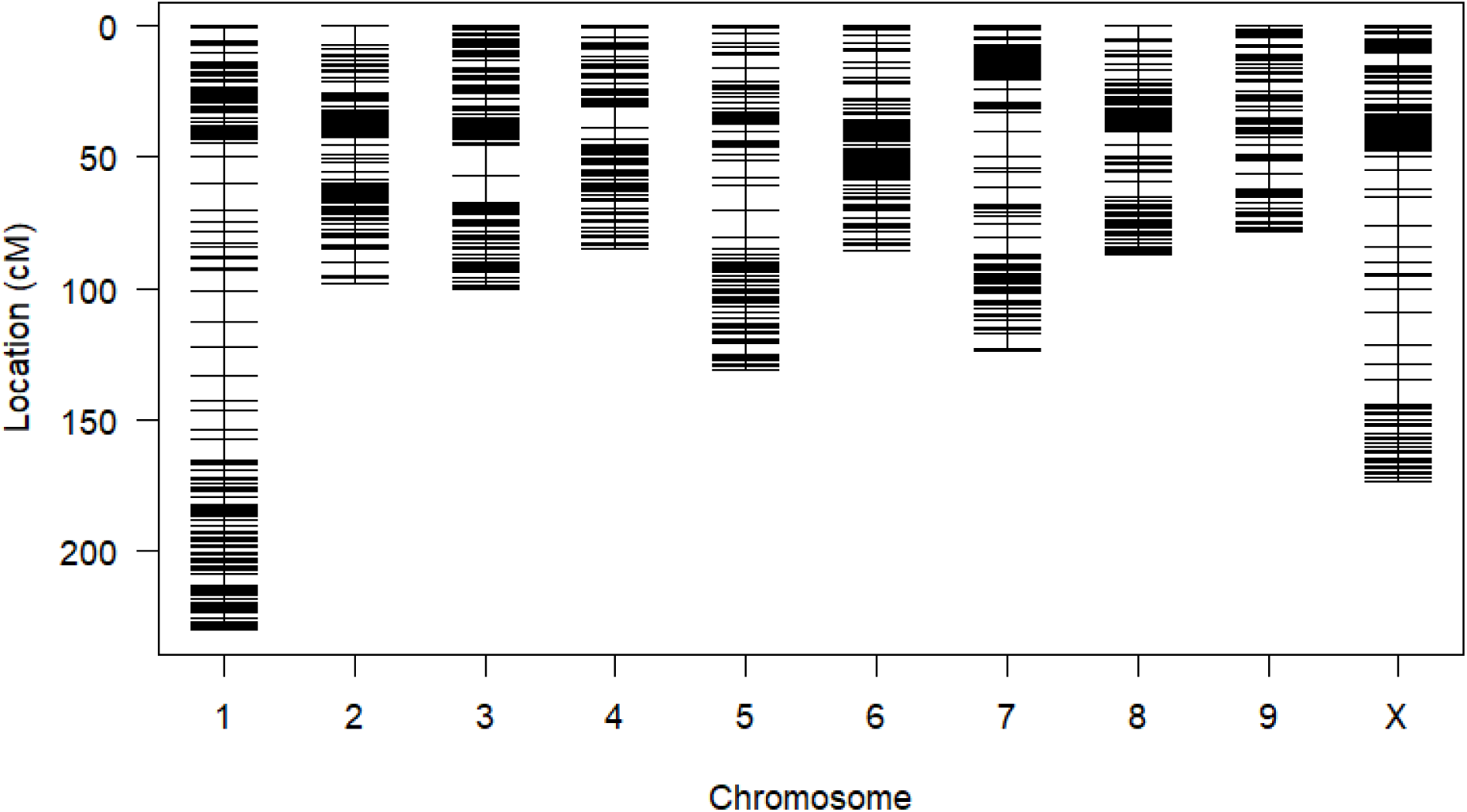
Genetic linkage map made using SNPs from an F_2_ population representing the 10 chromosomes of *Cannabis sativa*. Chromosome numbers are based on the numbering of the CBDRx reference genome (NCBI accession no: GCF_900626175.2).

**Table 3.** Summary statistics for linkage mapping in an F_2_ population of Cannabis sativa.

| Chromosome | Length<sup>a</sup> | Total Markers | Average Spacing<sup>a</sup> | Max Spacing<sup>a</sup> | % Coverage<sup>b</sup> |
| --- | --- | --- | --- | --- | --- |
| 1 | 230.4 | 174 | 1.3 | 11.5 | 99.50 |
| 2 | 98.4 | 109 | 0.9 | 7.5 | 99.95 |
| 3 | 100.4 | 121 | 0.8 | 11.4 | 99.67 |
| 4 | 84.6 | 94 | 0.9 | 8.6 | 99.82 |
| 5 | 130.8 | 101 | 1.3 | 10.1 | 99.94 |
| 6 | 85.8 | 104 | 0.8 | 5.7 | 99.90 |
| 7 | 124.0 | 116 | 1.1 | 9.2 | 98.04 |
| 8 | 87.4 | 87 | 1.0 | 5.7 | 92.31 |
| 9 | 78.6 | 81 | 1.0 | 5.6 | 99.98 |
| X | 174.0 | 116 | 1.5 | 12.7 | 100.0 |
| Overall | 1194.4 | 1103 | 1.1 | 12.7 | 98.91 |

<sup>a</sup>Length of linkage group, average spacing, and max spacing are all reported in centimorgan (cM).
<sup>b</sup>Percent coverage is based on the reference genome cs10 (NCBI Accession No: GCF\_900626175.2).

Flanking region sequence data for all markers retained in the linkage map were aligned to the ‘CBDRx-CS10’ reference genome. Of the 1,103 markers, 1,088 markers successfully aligned to the ‘CBDRx-CS10’ genome with 15 failures (Supplemental Table S2). Using the best alignment result for each marker, ‘Jamaican Lion’ contigs were assigned to ‘CBDRx-CS10’ chromosomes based on which chromosome a majority of a contig’s component markers aligned. Only 73 markers (6.7%) aligned to a different chromosome from the remainder of their contig (Figure 4), suggesting a high degree of collinearity in marker order between ‘CBDRx-CS10’, ‘Jamaican Lion’, and our assembled linkage map.

**Figure 4.**
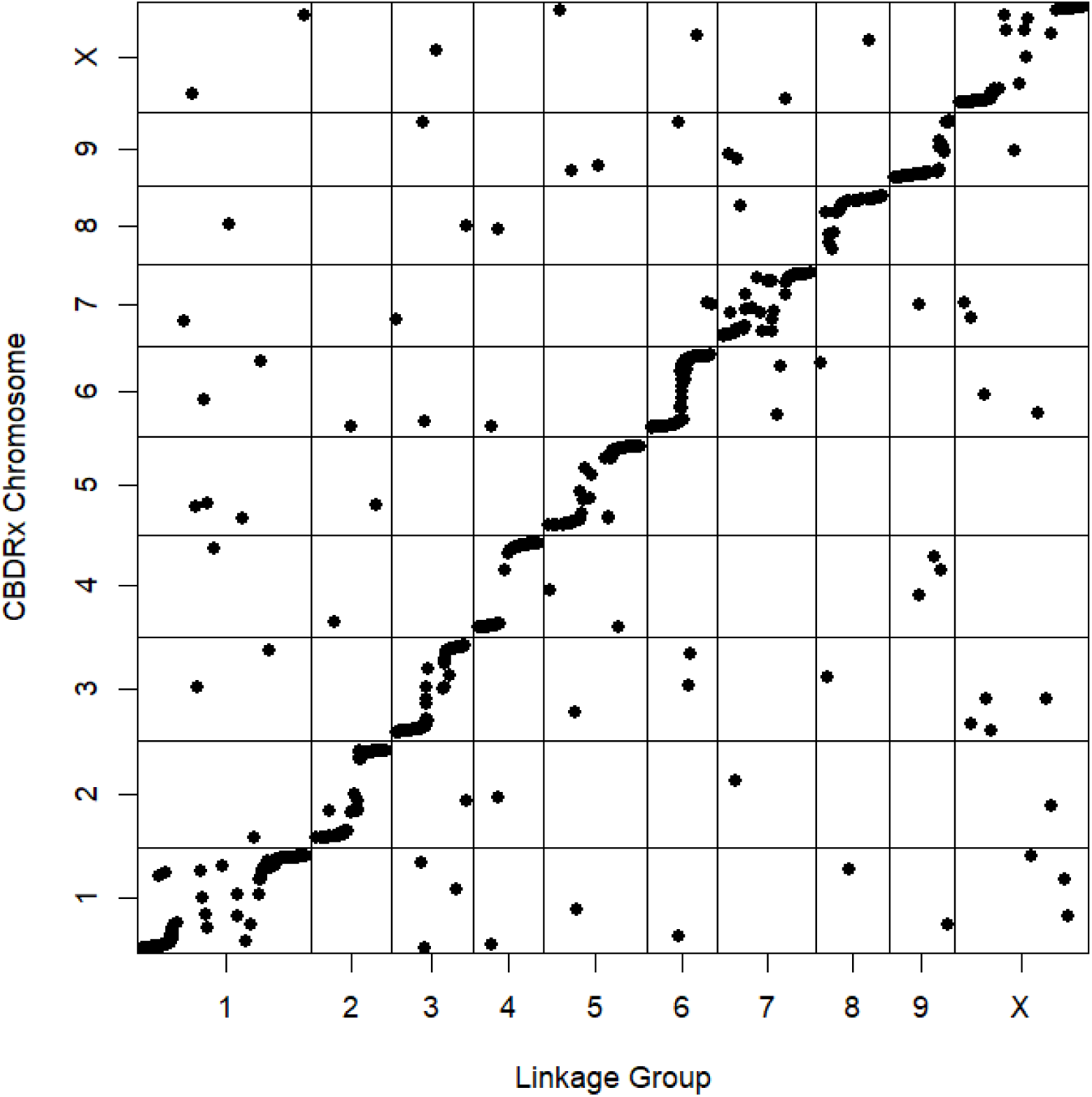
Graphical representation of the alignment of markers to the ‘CBDRx-CS10’ chromosomes as compared to linkage groups identified in the linkage mapping analysis. Alignments were conducted using BLASTn searches of a 1 Kbp region surrounding each SNP targeted by the CannSNP90 array.

Significant QTL mapped on both Chromosomes 1 and 4 had LOD scores of 47.8 and 11.2, respectively (Figure 5; Supplemental Table S2). The peak marker after QTL refinement on Chromosome 1 was contig856_3894427 while the peak marker on Chromosome 4 was contig7_81889. Although contig856_3894427 on Chromosome 1 is the peak, it carried missing data that was imputed as part of QTL mapping. Its close neighbor, contig504_3075882, was within the 1.5 LSI, with a similar LOD to the peak and a 100% call rate (Figure 5; Supplemental Table S2). Therefore, it is a more reliable representative marker for this QTL. Although contig504_3790889, previously identified using GAPIT as a perfect marker, is on the same ‘Jamaican Lion’ contig as our Chromosome 1 marker, it was not included in the linkage map due to its segregation as a backcross marker. It is worth noting that both markers from contig504 included in the linkage map were within the 1.5 LSI of the Chromosome 1 QTL, suggesting that this contig is highly associated with the day-neutral phenotype. While the QTL on Chromosome 4 was significant given its LOD score, the segregation pattern of the peak marker, contig7_81889, was unusual and imperfect. The day-neutral plants were, more often than not, called as heterozygotes which does not align with the conventional wisdom for this trait.

**Figure 5.**
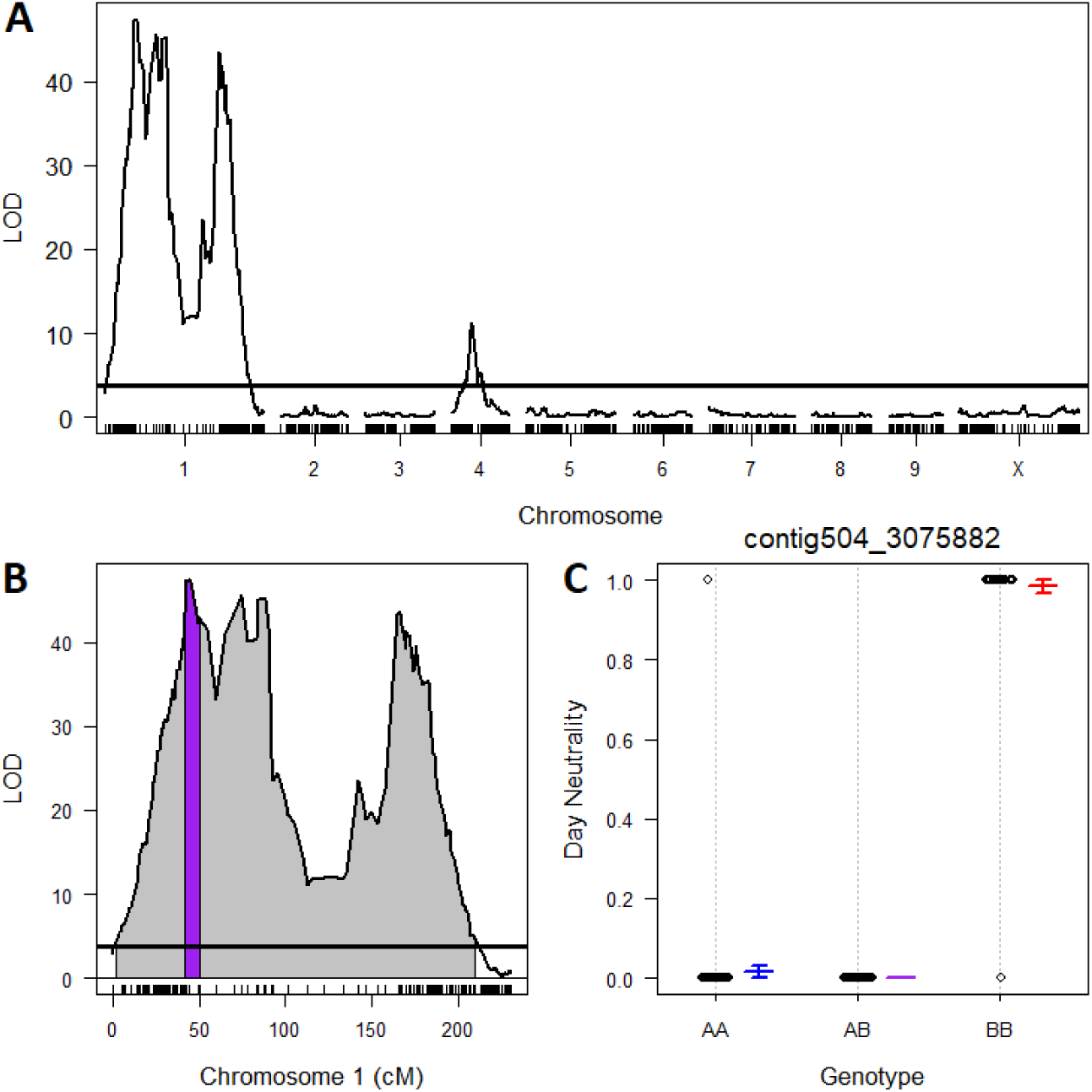
QTL analysis results. (A) QTL LOD scores for all ten chromosomes showing two QTL above our 5 percent permuted threshold (LOD = 3.69) associated with day-neutrality on chromosomes 1 and 4.)B) Focused LOD results for the QTL on chromosome 1. Grey area shows the region on chromosome 1 with a significant LOD score while the purple region is within the 1.5 LOD support interval for the peak marker, contig504_3075882. (C) Segregation of marker phase at contig504_3075882 in relation to day-neutrality.

Furthermore, several markers within the Chromosome 4 1.5 LSI had multiple BLAST hits outside of Chromosome 4 which could indicate either incorrect placement of these markers (although the only high similarity hit for contig7_81889 was to Chromosome 4), a biological difference in genome arrangement in our current crossing population and ‘CBDRx-CS10’, or a mis-assembly of contigs in the ‘CBDRx-CS10’ genome. Given the unlikelihood that there are two separate loci governing this trait, determined by the segregation patterns and previous literature surrounding this trait (Toth et al., 2022), this marker was not considered further.

### Physical location and candidate genes of the day-neutral trait

Using the MegaBLAST and BLASTn tools in NCBI, both loci identified by the GAPIT analysis, contig504_3790889 and contig262_2502527, can be placed on Chromosome 1 of *C. sativa* genotype CBDRx (GenBank accession no. LR213628.1), approximately 51 Mbp away from each other. Chromosome 1 is approximately 101 Mb in length in the ‘CBDRx-CS10’ assembly; locus contig504_3790889 BLAST hits place this marker near 23.7 Mbp and locus contig262_2502527 near 74.8 Mbp. BLAST searches for the SNP loci in the wild (non-cultivated) *C. sativa* genotype ‘JL’ place contig504_3790889 and contig262_2502527 on Chromosome 7 (GenBank accession no. CM022971.1) at approximately 19 Mbp and 54.2 Mbp, respectively. The discrepancies in chromosome numbers between ‘CBDRx-CS10’ and ‘JL’ are not likely indications of errors in WGS data, our SNP analyses, or BLAST searches. Rather, they indicate the lack of consensus and current shifting state of chromosome numbering in *C. sativa* that has yet to be fully resolved (Kovalchuk et al., 2020). In other words, Chromosome 1 of ‘CBDRx-CS10’ is syntenic with Chromosome 7 of ‘JL’. Markers contig856_3894427 and contig504_3075882 identified in the R/qtl analysis are located on Chromosome 1 at 26.8 and 21.4 Mbp, respectively.

While the exact locations of the markers identified in both the GAPIT and R/qtl analyses are unknown along Chromosome 1 in the F_2_ progeny, it is noteworthy that we have evidence of low levels of recombination between our key markers, contig504_3075882, contig504_3790889, and contig262_2502527, despite their apparent large physical distance. The refined QTL and 1.5 LSI for Chromosome 1 may cover a region of 9.6 Mb that encompasses contig504, however nearly the entirety of Chromosome 1 (0.64 - 99.64 Mb) bore a significant LOD score based on the initial permuted threshold covering contig262 (at 74.7 - 74.8 Mb), typically seen in regions with low recombination (Supplemental Figure S1) . Due to this large region of potential linkage to the day-neutral trait, recent patents have sought to protect gene editing across several potential gene candidates spanning the majority of Chromosome 1 (Phylos Biosciences, International Patent WO 2021/097496 A2). Previous studies in *C. sativa* have also suggested large regions of chromosomes that experience low rates of recombination, with higher rates of recombination happening only along the distal ends of the chromosomes (Kovalchuk et al., 2020).

The placement of the three markers, contig856_3894427, contig504_3075882, and contig504_3790889 are in close physical proximity to the *Autoflower1* region identified by Toth et al. (2022). The positions spanning marker contig856_3894427 are 17.6 to 27.2 Mbp; Toth et al. (2022) report a range between 17.74 and 22.94 Mbp. Toth et al. (2022) propose eight candidate genes in the region that have reported roles in flowering and development: *DOF ZINC FINGER NUCLEASES*, *NUCLEAR TRANSCRIPTION FACTOR Y SUBUNIT B-1* (*NFYB1*), *TARGET OF EARLY ACTIVATION TAGGED (TOE*)/*APETALA2* (*AP2*), *REGULATOR OF NONSENSE TRANSCRIPTS* 2/*UP-FRAMESHIFT 2* (*UPF2*), *ZINC FINGER CCCH DOMAIN-CONTAINING PROTEIN 11*, *PSEUDO-RESPONSE REGULATOR 3* (*PRR3*), *FAR1-RELATED SEQUENCE 5-LIKE*, and *LONG AFTER FAR RED 3* <u>(Toth et al. 2022)</u>. A patent submitted by Phylos Bioscience also names genes in the *AP2/TOE*, as well as *UPF2*, as a possible gene candidate for day-neutrality (International Patent WO 2021/097496 A2).

Leveraging whole genome sequencing (WGS) from a panel of three day-neutral recessive plants, we tested for variation in the proposed candidate genes. While day-neutral specific variation was found across the candidate proteins *PRR3*, *TOE/AP2*, and *UPF2*, both *PRR3* and *TOE/AP2* were predicted to have disruptions in key domains. *PRR3* was predicted to have a disrupted receiver (REC) domain (Figure 6A), while the second *AP2* domain has an insertion in *TOE/AP2* (Figure 6B). *PRR3* is known to impact photoperiodic flowering in two major crops, sorghum (*Sorghum biocolor*) and soybean (*Glycine max*) (Murphy et al. 2011; Li et al. 2019); *PRR3* is also a component of the core circadian clock that results in a circadian period length less than 24 hours (short period) when knocked out, and alterations in flowering time when overexpressed in *Arabidopsis* (Michael et al. 2003; Para et al. 2007). The REC domain is responsible for protein-protein interaction, while the CCT domain is responsible for DNA binding (Para et al. 2007; Gendron et al. 2012). The *PRR3* gene is missing from the *C. sativa* ‘Abacus’ genome assembly that was used for the Phylos Bioscience patent, possibly explaining why it was not identified as a candidate gene for day-neutrality in *C. sativa* (data not shown).

**Figure 6.**
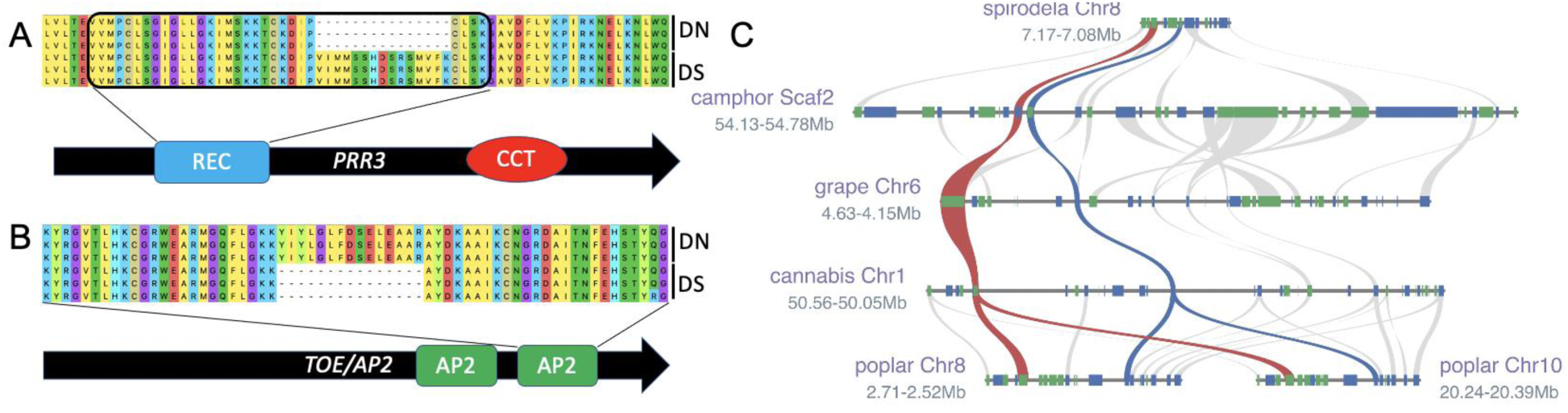
Day-neutral trait in an evolutionary conserved syntenic block that links phase change and photoperiodism to control flowering time. (A) *PSEUDO-RESPONSE REGULATOR 3* (*PRR3*) protein model in three day-neutral (DN) versus three day-sensitive (DS) plants reveals a knock-out of part of the receiver (REC) domain; the other DNA binding CCT (CONSTANS[CO], CO-like and TIMING OF CAB 1 [TOC1]) domain is not disrupted. (B) The day-neutral plants contained a predicted insertion in the *TARGET OF EARLY ACTIVATION TAGGED* (*TOE*)/*APETALA2* (AP2) DNA binding domain. (C) *PRR3* (red) and *TOE/AP2* (blue) are linked over evolutionary time from basal angiosperm like camphor (*Camphora officinarum*), to monocots (Spirodela polyrhiza), and across eudicots with different whole genome duplication (WGD) events like grape (*Vitis vinifera*) and cannabis (*Cannabis sativa*) that only has the Lambda triplication, and poplar (*Populus tremulus*) that has both the Lambda as well as a genus specific WGD. Gray lines represent other syntenic genes shared across the genomes.

*TOE/AP2* has also been implicated in flowering, but regarding its impact on phase transitions (Zhang et al. 2015; Werner, Bartrina, and Schmülling 2021; Luo, Yin, and He 2021; Du et al. 2020); loss of both *TOE1* and *TOE2* in *Arabidopsis* results in early flowering (Wu et al. 2009). Interestingly, *TOE/AP2* has also been implicated in trichome development (Wang et al. 2019). Since both *TOE/AP2* and *PRR3* proteins have day-neutral specific disruptions in key domains, it is possible that they both work together in a linked fashion to control flowering timing in *C. sativa*.

The result that there was low recombination in the region suggested that multiple genes in the region may play a role in the day-neutral phenotype. Recent work has shown that the organization of circadian, light signaling, and flowering time genes in the genome are under evolutionary constraints to remain linked in the genome (Michael 2022). Therefore, it was tested whether the genes in this region were also under evolutionary constraint and inherited together over time, which could mean they work together to impact the day-neutral flowering trait.

Leveraging a dataset that determined the evolutionary trajectory of all genes across 123 high quality plant genomes (Zhao and Schranz 2019; Zhao et al. 2021), we found that in 104 (85%) plant genomes spanning basal eudicots to monocots *PRR3* and *AP2/TOE* are inherited in larger than normal syntenic blocks (Figure 6C). In many species, including *Arabidopsis*, these genes are directly next to one another, which is consistent with functionally linked genes moving closer together after more rounds of whole genome duplication (WGD) and fractionation like was noted for other circadian, light signaling, and flowering time genes (Michael 2022). In contrast to the linkages between other circadian clock genes (*CCA1-PRR9*, *RVE4-PRR7*, and *PIF3-PHYA*), which do not have any linkages in the monocot grasses, there are seven grasses including rice (*Oryza sativa*) and setaria (*Setaria* spp.) that have *PRR3* and *TOE* linked in a syntenic block in their genome.

This evolutionary linkage suggests that *TOE/AP2* and *PRR3* may work together to control flowering. Leveraging the framework outlined by Spitzer-Rimon et al. (2019) proposing two stages of flowering in *C. sativa* (described in the Introduction), it is possible that *TOE/AP2* and *PRR3* are playing distinct roles the day-neutral phenotype; *TOE/AP2* may be monitoring plant age to enable the plant to phase transition to a permissive flowering state, while *PRR3* may be integrating the photoperiodic information with the circadian clock leading to the terminal flower and bulking. These results support the growing evidence that functionally relevant gene neighborhoods are more prevalent in plants; testing this hypothesis would require gene editing technologies such as CRISPR.

### Significance of findings

While the specific markers used in this study are likely to have limited value in other crosses of *C. sativa*, the information presented here, including flanking region sequence data, the physical location of these markers in publicly available genomes, and the variation in candidate genes can be used by breeders and researchers to re-design markers as well as editing strategies to further elucidate this important trait.

## ACKNOWLEDGMENTS

We would like to thank Kevin McKernan of Medicinal Genomics for sharing flanking region sequence data for the purposes of developing our qPCR markers. This work was partially supported by the New York State Department of Agriculture and Markets through grants (AC477 and AC483) to L. B. S. from Empire State Development Corporation as well as funding to L. B. S. through a cooperative research agreement with the United States Department of Agriculture Agricultural Research Service.

## SUPPLEMENTAL MATERIAL

Supplemental materials include Supplemental Tables S1 and S2 which describe the additional cultivars tested with the markers developed in this study and the BLASTn results for flanking region sequence data, respectively. Supplemental Figure S1 shows pairwise recombination frequencies and LOD scores among markers identified during QTL analysis.

Supplementary data of additional sequence data for the parents of the F_1_ parent are also included.

## REFERENCES

Beutler, J. & Der Marderosian, A. (1978). Chemotaxonomy of *Cannabis* I. Crossbreeding between *Cannabis sativa* and *C. ruderalis*, with analysis of cannabinoid content. Economic Botany, 32, 378–394.

Broman K. W., Wu, H., Sen, Ś., & Churchill, G. A. (2003). R/qtl: QTL mapping in experimental crosses. Bioinformatics, 19(7), 889–890.

Cascini, F., Farcomeni, A., Migliorini, D., Baldassarri, L., Boschi, I., Martello, S., Amaducci, S., Lucini, L., & Bernardi, J. (2019). Highly predictive genetic markers distinguish drug-type from fiber-type *Cannabis* sativa L. Plants, 8, 496.

Compacho, C., Coulouris, G., Avagjan, V., Ma, N., Papadopoulos, J., Bealer, K., & Madden, T. L. (2009). BLAST+: architecture and applications. BMC Bioinformatics, 10, 421.

de Meijer, E. P., Bagatta, M. Carboni, A., Crucitti, P., Moliterni, V. C., Ranalli, P., & Mandolino, G. (2003). The inheritance of chemical phenotype in *Cannabis sativa* L. Genetics, 163, 335–346.

Du, S-S., Li, L., Li, L., Wei, X., Xu, F., Xu, P., Wang, W., Xu, P., Cao, X., Miao, L., Guo, T., Wang, S., Mao, Z., & Yang, H-Q. (2020). Photoexcited cryptochrome2 interacts directly with *TOE1* and *TOE2* in flowering regulation. Plant Physiology, 184, 487–505.

Garfinkel, A. R., Otten, M., & Crawford, S. (2021). SNP in Potentially Defunct Tetrahydrocannabinolic Acid Synthase is a Marker for Cannabigerolic Acid Dominance in *Cannabis sativa* L. Genes, 12(2), 228.

Gendron, J.M., Pruneda-Paz, J.L., Doherty, C.J., Gross, A.M., Kang, S.E., & Kay, S.A. (2012). *Arabidopsis* circadian clock protein, *TOC1*, is a DNA-binding transcription factor. PNAS, 109, 3167–3172.

Hall, J., Bhattarai, S. P., & Midmore, D. J. (2012). Review of flowering control in industrial hemp. Journal of Natural Fibers, 9, 23–36.

Hanuš, L.O., Meyer, S. M., Muñoz, E., Taglialatela-Scafati, O., & Appendino, G. (2016). Phytocannabinoids: a unified critical inventory. Natural Product Reports, 33, 1357–1392.

Henry, P., Khatodia, S., Kapoor, K., Gonzales, B., Middleton, A., Hong, K., Hilyard, A., Johnson, S., Allen, D., Chester, Z., Jin, D., Rodriguez Jule, J. C., Wilson, I., Gangola, M., Broome, J., Caplan, D., Adhikary, D., Deyholos, M. K., Morgan, M., Hall, O. W., Guppy, B. J., & Orser, C. (2020). A single nucleotide polymorphism assay sheds light on the extent and distribution of genetic diversity, population structure and functional basis of key traits in cultivated north American cannabis. Journal of Cannabis Research, 2(1), 26.

Lewis, M. A., Russo, E. B., & Smith, K. M. (2018). Pharmacological foundations of *Cannabis* chemovars. Planta Medica, 84, 225–233.

Li, M-W., Liu, W., Lam, H-M., & Gendron, J.M. (2019). Characterization of two growth period QTLs reveals modification of *PRR3* genes during soybean domestication. Plant & Cell Physiology, 60, 407–20.

Lipka, A.E., Tian, F., Wang, Q., Peiffer, J., Li, M., Bradbury, P. J., Gore, M. A., Buckler, E. S., & Zhang, Z. (2012). GAPIT: genome association and prediction integrated tool. Bioinformatics, 28, 2397–2399.

Luo, X., Yin, M., & He, Y. (2021). Molecular genetic understanding of photoperiodic regulation of flowering time in *Arabidopsis* and soybean. International Journal of Molecular Sciences, 23. https://doi.org/10.3390/ijms23010466.

Mahlberg, P. G. & Kim, E. S. (2004). Accumulation of cannabinoids in glandular trichomes of *Cannabi*s (Cannabaceae). Journal of Industrial Hemp, 9, 15–36.

McPartland, J. M. (2018). *Cannabis* systematics at the levels of family, genus, and species. Cannabis and Cannabinoid Research, 3, 203–212.

Michael, T.P. (2022). Core circadian clock and light signaling genes brought into genetic linkage across the green lineage. Plant Physiology, 190, 1037–56.

Michael, T.P., Salomé, P.A., Yu, H.J., Spencer, T.R., Sharp, E.L., McPeek, M.A., Alonso, J.M., Ecker, J.R., & McClung, C.R. (2003). Enhanced fitness conferred by naturally occurring variation in the circadian clock. Science, 302, 1049–1053.

Murphy, R.L., Robert, R. Klein, D.T. Morishige, J.A. Brady, W.L. Rooney, Miller, F.R., Dugas, D.V., Klein, P.E., & Mullet, J.E. (2011). Coincident light and clock regulation of pseudoresponse regulator protein 37 (*PRR37*) controls photoperiodic flowering in sorghum. PNAS, 108, 16469–16474.

Onofri, C., & Mandolino, G. (2017). Genomics and Molecular Markers in Cannabis sativa L. In Cannabis sativa L.-botany and biotechnology (pp. 319–342). Springer, Cham.

Para, A., Farré, E.M., Imaizumi, T., Pruneda-Paz, J.L., Harmon, F.G., & Kay, S.A. (2007). *PRR3* is a vascular regulator of *TOC1* stability in the *Arabidopsis* circadian clock. The Plant Cell, 19, 3462–3473.

Piunno, K. F., Golenia, G., Boudko, E. A., Downey, C., & Jones, A. M. P. (2019). Regeneration of shoots from immature and mature inflorescences of *Cannabis sativa*. Canadian Journal of Plant Science, 99, 556–559.

R Core Team (2022). R: A language and environment for statistical computing. R Foundation for Statistical Computing, Vienna, Austria. URL https://www.R-project.org/.

Ram, H. M. & Sett, R. (1982). Induction of fertile male flowers in genetically female *Cannabis* sativa plants by silver nitrate and silver thiosulphate anionic complex. Theoretical Applied Genetics, 62, 369–375.

Small, E. (2018). Dwarf germplasm: the key to giant *Cannabis* hempseed and cannabinoid crops. Genetic Resources in Crop Evolution, 65, 1071–1107.

Spitzer-Rimon, B., Duchin, S., Bernstein, N., & Kamenetsky, R. (2019). Architecture and florogenesis in female *Cannabis sativa* plants. Frontiers in Plant Science, 10, 350.

Taylor J. & Butler, D. (2017). R Package ASMap: efficient genetic linkage map construction and diagnosis. Journal of Statistical Software, 79(6), 1–29.

Toth, J. A., Stack, G. M., Cala, A. R., Carlson, C. H., Wilk, R. L., Crawford, J. L., Viands, D. R., Philippe, G., Smart, C. D., & Rose, J. K. (2020). Development and validation of genetic markers for sex and cannabinoid chemotype in *Cannabis sativa* L. GCB Bioenergy, 12, 213–222.

Walia, K. (2019). Global Cannabidiol market size, share, trends. Industry Report, 2026. 21 November 2019. https://works.bepress.com/kritika-walia/260/.

Wang, L., Zhou, C-M., Mai, Y-X., Li, L-Z., Gao, J., Shang, G-D., Lian, H., Han, L., Zhang, T-Q., Tang, H-B., Ren, H., Wang, F-X., Wu, L-Y., Liu, X-L., Wang, C-S., Chen, E-R., Zhang, X-N., Liu, X-N., Liu, C., Wang, J-W. (2019). A spatiotemporally regulated transcriptional complex underlies heteroblastic development of leaf hairs in *Arabidopsis thaliana*. The EMBO Journal, 38. https://doi.org/10.15252/embj.2018100063.

Werner, S., Bartrina, I., & Schmülling, T. (2021). Cytokinin regulates vegetative phase change in *Arabidopsis thaliana* through the *miR172/TOE1-TOE2* module. Nature Communications, 12, 5816.

Wu, G., Park, M.Y., Conway, S.R., Wang, J-W., Weigel, D., & Poethig, R.S. (2009). The sequential action of miR156 and miR172 regulates developmental timing in *Arabidopsis*. Cell, 138, 750–59.

Yang, R., Berthold, E., McCurdy, C.R., da Silva Benevenute, S., Brym, Z. T., & Freeman, J. H. (2020). Development of cannabinoids in flowers of industrial hemp (*Cannabis sativa* L.): a pilot study. Journal of Agricultural and Food Chemistry, 68(22), 6058–6064.

Zhang, M., Anderson, S. L., Brym, Z. T., & Pearson, B. J. (2021). Photoperiodic flowering response of essential oil, grain, and fiber hemp (*Cannabis sativa* L.) cultivars. Frontiers in Plant Science, 1498.

Zhang, B., Wang, L., Zeng, L., Zhang, C., & Ma, H. (2015). *Arabidopsis TOE* proteins convey a photoperiodic signal to antagonize *CONSTANS* and regulate flowering time. Genes & Development, 29, 975–987.

Zhao, T. & Schranz, E.M. (2019). Network-based microsynteny analysis identifies major differences and genomic outliers in mammalian and angiosperm genomes. PNAS, 116, 2165–2174.

Zhao, T., Zwaenepoel, A., Xue, J-Y., Kao, S-M., Li, Z., Schranz, M.E., & Van de Peer, Y. (2021). Whole-genome microsynteny-based phylogeny of angiosperms. Nature Communications, 12, 3498.

