## Supplemental Data for "Genetic Mapping of SNP Markers and Candidate Genes Associated with Day-Neutral Flowering in *Cannabis sativa* L"

This document contains supplemental material from:

No. Pages: 5
No. Figures: 1
No. Tables: 1
Supplementary Sequence Data supplied on pages 5-6.

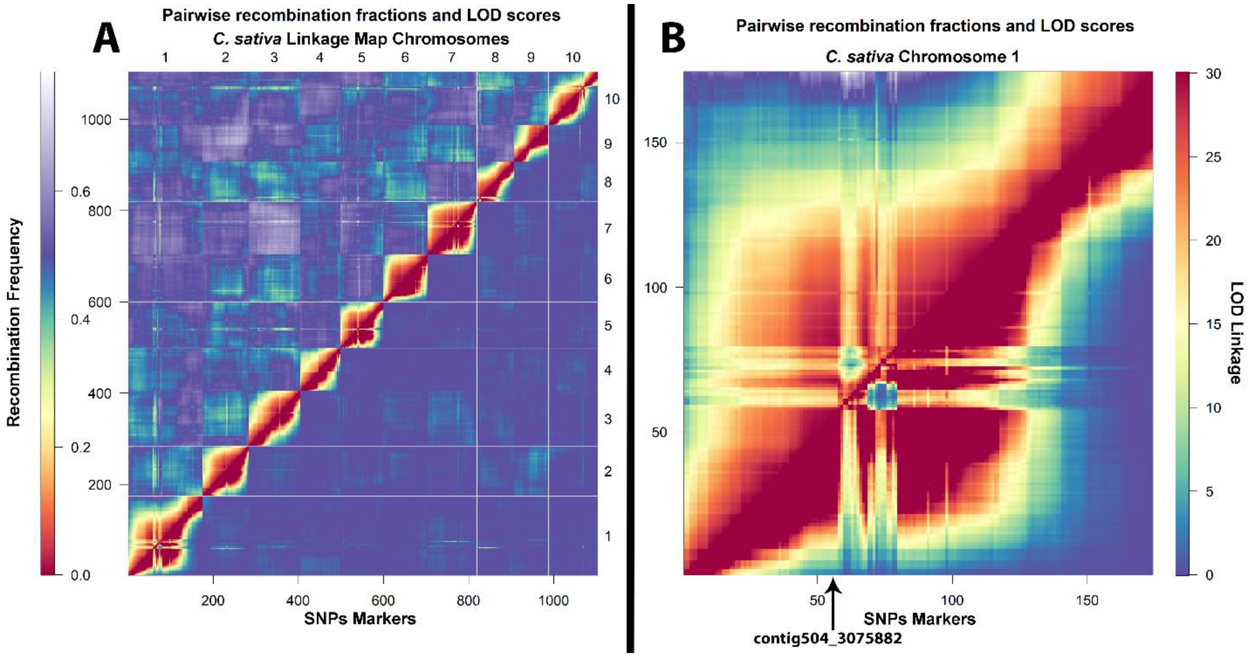


Supplemental Figure S1. Pairwise recombination frequencies and LOD scores. Visualizes low recombination and therefore strong linkage within and throughout each linkage group for (A) all chromosomes within the linkage map and (B) specifically chromosome 1. The arrow in panel B designates the position of contig504_3075882, the representative marker identified through QTL mapping.

Supplemental Table S1. Genotypes of non-relative *Cannabis sativa* cultivars at two SNPs that were identified to be associated with the day-neutral phenotype in an $F_2$ crossing population.

| Cultivar | Seed Breeder | Phenotype[a] | SNP_504[b] | SNP_262[bc] |
|---|---|---|---|---|
| TS1-3 | Oregon CBD | PS | AA | TT |
| Sour Tsu | Larry Ringo | PS | AA | TC |
| Charlotte's Web | Stanley Brothers | PS | AA | TC |
| Suver 8 | Oregon CBD | PS | AA | CC |
| W19 | Oregon CBD | PS | AA | TC |
| SS | Oregon CBD | PS | AA | CC |
| AcDc (phenotype of 'Cannatonic') | Resin Seeds | PS | AA | TC |
| DC Haze CC | Oregon CBD | PS | AA | TT |
| Dinamed Kush | Dinafem Seeds | DN | AA | TT |
| OG Kush | Dinafem Seeds | DN | AA | TC |
| Grapey Walter | Mephisto Genetics | DN | AA | TT |
| White Crack | Mephisto Genetics | DN | AA | TC |
| Crème De La Chem | Mephisto Genetics | DN | AA | TT |
| Forum Stomper | Mephisto Genetics | DN | AA | CC |
| Samsquanch OG | Mephisto Genetics | DN | AA | TT |
| Sour Crack | Mephisto Genetics | DN | AA | TC |
| Alien Vs. Triangle $F_2$ | Mephisto Genetics | DN | AA | TT |

[a]DN = Day-neutral; PS = Photoperiod-sensitive

[b]qPCR markers identifying the former two loci as markers for the DN trait.

[c]Genotypes in gray represent genotype calls which do not match the phenotype.

Supplemental Table S2. BLASTn results for flanking region sequence data is an Excel file uploaded separately.

Supplemental Sequence Data. Flanking region sequences from ‘ERB’ with the SNP targeted in qPCR analyses identified.

>ERB_SNP_contig262_000012F|arrow
TACTAAATCTAATATAACAATATTACTAGCTCATTATTTAGTACTTATTTGCTTCCTC
TCTAATTTCCTTTCTTTCTAAGTACAATCCACCTTTCTCAAATTACAATGGTAACTCTT
TCTTTTTCTTACTATTTTCTTTTCATTCATGTTCATCAACACATTAATACATAAATATG
TACATATTGGTTTTCATCATTTGTAATTTGAAGATAGTTTTTTAAAATTTTCTTGTTGG
ATAGTTTCGCCTAATTTCCTCTATATTTGTTGTTATATGAAAGGAGTTGAAGAGATAT
CTTTGCAAAGCCACACATATTTTTTTTCCTAACAAGACAGATTTTCTCATATATATAT
ATATACATATATAATATACCATTACTTTTCTCTTTTAGTAGTTCCCCAACCTTAAAAC
GAACATGATGTTAAAAACACAGATGATAACTCTAAAAAACAATTGGAAAATTTTGT
GCCAGAAAGTAAAAGCTCATTCTCATGCCTTTATTTTCAGGTAAAAGCAATTTCTTTC
TAACTCAAACTCTTCTCTTTCCCTTTTCTTTAACAGCGTTGTATTACTTTTCTGTTGTG
TGATCAGGACAGTGAGAACAAAGGTAGCTATGAGGAAAGCTTTGATCAAGAAGCTT
TGCCACTCAAAACCGCATTTGAGTTATTTGATGCCAACTTTTTCAGTAGTAAAAAGG
TCACAAATTTAATCAATTATATGGTTAATAATACACTAGCTAATCATTACTTCTCATT
CTATATCTTTGGCTATTTATAGATGGTGGAAATTACTAAGGGTGCAAAAGAGCTCAA
CGTCCCTACTATTAGAGCAAATAGAAAGCTTGTTGGTTCTGTTAATGGAGGGCTTTA
TCATCATTCTCCTTTGGTCTTCAACCCTGAATGGGCAACTGAAGAAGCTAACAACAA
AAAAAAGAGATTTAATTATCCTT[A/G]TTTGGTTGGCACACAAAGACCAAACAAAGA
AGATGATATTGCCTTCATGAGTGTAATTAAGAAAAGATTCTTTGATATTTTTCATATC
TACATGTGTGTGCTGCTGTGATAATAAGTAAACCCATTTGATGATTTTCAGATTCTTG
AACTAGGGGAGCTGATTAAGACAAAGCAAGTTACATCTGAGGAGCTCACTCAAATT
TGTCTTAAAAGACTGAAAAGGTAATGAAAAAAGTCCATATTTTGTTCTAATTCAAGG
CTCCCTTTTTTGACTGATGTAGTAATTGTGTGTAGGTATAATCCTGTTCTTGAGGCTG
TTATCTCTTACACTGAAGACTTGGCATACAAGCAAGCAAAAGAAGCTGATGAATTGT
TCTCCAGAGGAGTGTACTTGGGTATAGACCTCTTTTTCCACTAACTCATTAAGTAAAT
ATTTACTTTATGGTTTTTATATAATAGAAAGTTCTTCATTACAATGCTTGAATTAGGT
CCTCTCCATGGGATCCCTTATGGGTTAAAGGATATAATTTCAGTACCCCAATACAAA
ACAACATGGGGTTCAAAAAGTTTCAAAGATCAAGTCCTTGACATTGAAGCTTGGGTT
TACAAAAGGTAACAAATATGTTCTGAATATAGAAGTTGCATCTATAGAACATTGATA
TTCTAGTTTATGACTGTACATATGTGTTGAATGAAGGTTGAAGTCTGCTGGGGCAGT
TCTTGTAGCCAAACTTGTTTCTGGATCACTGGCATATGATGATATATGGTTTGGTGGT
AGGACAAGGAACCCTTGGAATATTGAAGAATTCTCCACAGGTTCTTCAGCTGGCCCA
GCTGCCTGCATCTCAGCTGGTATTATTTTTGAGCTTCTTGGTTTTCTTTTTTTCAATTA
ATATATGGTTGTCTTGACTAGTAACTTTGTATGTGTGAACTGAAAAATCACAACTTC
ACTTGGATCAAGAATGGTTCTCTCAGATTTTTTTCTTTTAACAGGGTTGGTTCCATTT
GCTATTGGATCAGAAACAGTTGGTTCGATGACCTTGCCTGCTTCGC

>ERB_SNP_504_000014F|arrow
ATGCCAATACATAATATTTGACTTTTCAATTTGATCCATCAAGGCATCAAATATATGT
ATTGATCCAAGGAGCCATTAATCACAAGCCAAGCGTTTTTCTTCACCCATTTTGAATT
TGTACCTGGGATTTTCTTCTTGGCGGTAATTTAATGGAAATCAAAATTTATTATTATA
TATTATTGTGTAAGCGTTAGTCGAAAATGGGTTTTCTTTTACAGTAGTAGTACTACTC
AAGCTTTGATAAGTTCAGCACATTTAATGGAATTTGTTATTGTTGTCTTGTTGATAAG
TTCAGGTTCAACACATATATCATTGTAATGTTATGAGGTTTTTAGAGTGTGAAATCAT
CTTCTTATGTTATAATGATACTTACTGGTTGTTCTTTTCAATGATTGTAATTTGTATAT
GCCAAAAAAAATGTTTCATGATTTGGTCTGTGTTGCTGGTTTTATTGTTTTATTTTGTT
TTTGTGAAACTTTATCTATTCCTATGGGAGAAGTCCATAAATGATGCTTTACAGCTTT
TGATTGAAATTATTGAGTTCTATATATGTACATATAAGTAAGTAATTAATCTTTTTAT
GTTTCCTTTTTTTTTTTTCATATATATATATATATATACATATTTAATTGGTTTATATA
AACGTGCATATTCAATTTTTGGTTAGAATTTTATTTCTTAATGATTCGCTTGTATTTCT
ATATAGCTTCATCTTCATGGGATAGTTATTGAAATTTTTGTTTGATTTTTCAGAAACT
GAGAAGAAGCTTAGGAACTCAGAAGGGTTAATTTTAATGTGTTAGATGGTGCAAAT
GAGTTAAGGTGAGGACAATTTTATTATACTAGCGTTACAGAAATGGAAGGTAAGCA
AAGTGTCATTGCAAGACTAATGGGCTTTGATGAACACCAACCTCTGGAAGCTATCCA
TAAGCCACAGAGAGTACTCTCTGAGAATTATTTGAGGAAAGTTGCTTCTATTGGTGT
TGGGAAGAAGAAGAA[T/G]AAGAAGGAAGCCAAGGAGGTTTGTCAAAGAAAAGGA
GTCTTGGATTTGTCATTTCAAAAATGTGCAACTGGAGAAGAGTCTTCAAAGTTGGAT
TTTTTTCAAAAACATAAGAATGACAGTGTCACATGTTCTGAGAGATTTGACTTGGTA
AGACAATATGAATCAAGTTCTTCGCATATGAGAATTCTTGTTTTGAAACCAATATCC
GGGAAGGCTAAGAGATTTGGAGTAAAGAATGTTCAGTGCAGTGTGGAATCTATTAG
ACAAAGATCTTCTGCCTCAAGGGAAATCTTTAAGGAGCTTAAAAGAATAAGAAGAT
GGAGGAATTTAGGCTTTCCAACAATGAACTCAAGCTTAGGATTAAGTGGTGGCGAC
ACTCCTACATTAAAATCTGAGATTTTGTCACCTCCTTCCACTTGTTCTTTTGAGTACT
CAGATAAATCAACTTTAAGCAACAGATCCAAAAAGCAGATTTGGGAAAGGTGTCTG
TTGAGTGAAAAATATAAAGTTGAAACGACTTCTAGAATTAGTACCCCAGATGAGCT
GATTGCTTTGCAAGATCCTAATTCATGGCCTAGAAATTGCAAGCCTGGTCTGGACCT
CAGTAAGTTTGCCAAACAACAACTCAAGTGTGAAGCTTATAGCAACAGATGGTATTT
GGGGCTAGAAAAGCCCCTTAATCAGGCAAGGAGTAAGATGATAATAAAACATGATT
TTATAAAAGATATGGTATTAGACAAAGAGATTGAGAACACAAATAAATCCTCTGGA
GAATCTGAGACTGAGAGTGAGCAATTAGACCCTACCATGTGCATCAGCTTAGAAGA
AGACTATGATTCTTCTAGCCTAACTACAGATACTTCAACTCAACAGGTTTGTTTCTTT
TTTCATCATAAGAACACTTTTTTCTAAAGGATATTATTGGCTTACTTAACTATGAAGT
TAGGATTCCTCTTTCTATGAGTCTTGTTAACCTCAACTTAGGAAAGTGATGCTTCAAA
TGTAT
